# Single VLP lipid-mixing measurements confirm off-pathway state in dengue virus fusion mechanism

**DOI:** 10.1101/2025.03.21.644571

**Authors:** Tasnim K. Anika, Fiona Campbell, Bianca Linden, Connor J. Criswell, Miranda Kimm, Priscilla Li-ning Yang, Robert J. Rawle

## Abstract

Dengue virus (DENV) is the causative agent of dengue fever and exerts a substantial healthcare burden worldwide. Like other flaviviruses, DENV must undergo membrane fusion with the host cell in order to initiate infection. This membrane fusion occurs following acidification during endocytosis and is pH dependent. Here, we interrogate whether the mechanism of DENV fusion contains an off-pathway state, such has been reported previously for two other flaviviruses - Zika virus and West Nile virus. To do this, we utilize single particle lipid mixing measurements of DENV virus-like particles (VLPs) to tethered liposomes, together with computational modeling inspired by chemical kinetics. By observing and then modeling the pH dependence of single VLP fusion kinetics, we provide evidence that the DENV fusion mechanism must contain an off-pathway state. Measuring the proportion of VLPs undergoing hemi-fusion over time, we also demonstrate that the off-pathway state appears to be slowly reversible over tens of minutes, at least for some virions. Additionally, we find that late endosomal anionic lipids do not appear to influence the off-pathway mechanism to any great extent. In conjunction with the prior reports on Zika virus and West Nile virus, this work indicates that an off-pathway fusion state may be a feature of flavivirus fusion more broadly. We also note that the platform and mechanistic model described in this study may be useful in elucidating the mechanism of action of small molecule inhibitors of flavivirus fusion developed by our group and others.

**Statement of Significance:** Dengue virus (DENV) causes dengue fever and infects an estimated hundreds of millions of people annually. To date, there are no specific antiviral drugs for DENV and limited vaccination options, highlighting the need to better understand this important pathogen. In this report, we investigate the mechanism of DENV membrane fusion, an early step in the viral infectious cycle, using a mix of experimental techniques and computer simulations. We find strong evidence that the DENV fusion mechanism contains an off-pathway state, in which it can get stalled prior to membrane fusion. Understanding this off-pathway state could be an avenue to develop antiviral strategies against DENV and other related viruses.

## 1. Introduction

Dengue virus (DENV), the causative agent of dengue fever, is a membrane-enveloped, positive-sense RNA virus and a member of the *Flaviviridae* family. It is primarily transmitted to humans by mosquitoes and the numbers of annual infections worldwide are estimated to be in the hundreds of millions (1). While many dengue infections result in asymptomatic or mild disease, severe dengue occurs with some regularity, and can lead to hospitalization and death (2). To date, there are no antivirals specific for DENV, and while there is currently one approved vaccine (QDenga), it is limited to children in high risk geographical areas (2). This underscores the need for ongoing research into this important pathogen.

Like other flaviviruses, the initial stages of DENV infection are first binding to the host cell membrane, followed by internalization via clathrin-mediated endocytosis, whereupon membrane fusion occurs between the viral and endocytic membranes (3). Fusion is mediated by the transmembrane viral E protein, a class II fusion protein which exists in close-packed dimer pairs on the viral membrane surface (4). Upon exposure to low pH in the endosome, the E protein is activated, extending and inserting a hydrophobic fusion loop into the host membrane, and arranging into trimers (5). These extended trimers are unstable and ultimately re-fold, bringing the viral and endocytic membranes into close apposition, leading to membrane fusion (6).

In recent years, researchers studying viral fusion of many viral families have begun employing sensitive fluorescence microscopy-based measurements of individual viruses or viruslike particles (VLPs) fusing to self-assembled model membranes (7–15). We and others have utilized this approach to investigate the mechanism of flavivirus fusion (16–18), and have examined the pH-dependence of the fusion of West Nile virus (WNV) and Zika virus (ZIKV), two other prominent members of the *Flaviviridae* family. In those studies, it was observed that the distribution of wait times between low pH exposure and membrane fusion (observed as lipid mixing with the target membrane) was remarkably insensitive to pH. On the other hand, the viral fusion extent (number of viral particles fused during the observation window/number observed) decreased substantially with increasing pH. Computational modeling of these data indicated that in order to adequately model both observations, the fusion mechanism must contain an off-pathway state in which the virus is aborted from productive fusion (16, 19). This was in contrast to prior work, which postulated that a linear model would best describe the flavivirus fusion process (17), similar to linear fusion mechanisms utilized by other viral families (7, 9).

In this report, we examine whether this off-pathway state may be broadly characteristic of mosquito-borne flaviviruses. To do this, we adapted previous single virus fusion assays to study the pH-dependence of DENV fusion. Then, using similar computational modeling approaches as in prior reports, we investigated whether the fusion mechanism of DENV-2 also possesses an off-pathway state, similar to WNV and ZIKV. We also performed experiments to examine the reversibility of the off-pathway state, as well as its dependence on the lipid composition of the target membrane.

## 2. Results/Discussion

### 2.1. Adaptation of single virus fusion assay to study DENV VLPs

To study the fusion of dengue virus, we adapted a single viruslike particle (VLP) lipid mixing assay (see schematic in **Figure 1a**) that has previously been used to study a number of other viruses (10, 12, 16, 20–22). In brief, liposomes (∼120-130 nm in diameter as determined by DLS) were tethered via NeutrAvidin-biotin interactions to a glass coverslip coated with PLL-PEG-biotin inside a PDMS microfluidic flow cell. These liposomes served as the target membranes for fusion. DENV VLPs were produced in HEK293 cells and were characterized by negative stain electron microscopy as uniform particles ∼20-40 nm in diameter (**Figure 1c**). Texas Red-DHPE was used to label the lipid envelope of the VLPs at a self-quenched concentration. Complementary DNA-lipids were incorporated into the labeled VLPs and target liposomes respectively (see Materials and Methods), enabling the VLPs to bind to the tethered liposomes by DNA hybridization. This DNA-lipid approach to facilitate virus-target binding in the absence of a viral receptor has been validated previously for other viruses, including influenza, Zika, and SARS-CoV-2 viruses (10, 12, 16, 20, 23). Unbound VLPs were removed by buffer rinsing, and fusion of the bound VLPs was triggered by introduction of a low pH buffer into the flow cell. Fusion was observed via fluorescence video microscopy as fluorescence de-quenching of the Texas Red-DHPE upon lipid mixing between the VLP and tethered liposome. The waiting time between pH drop and lipid mixing was then determined (**Figure 1b**) for each fusion event.

**Figure 1.**
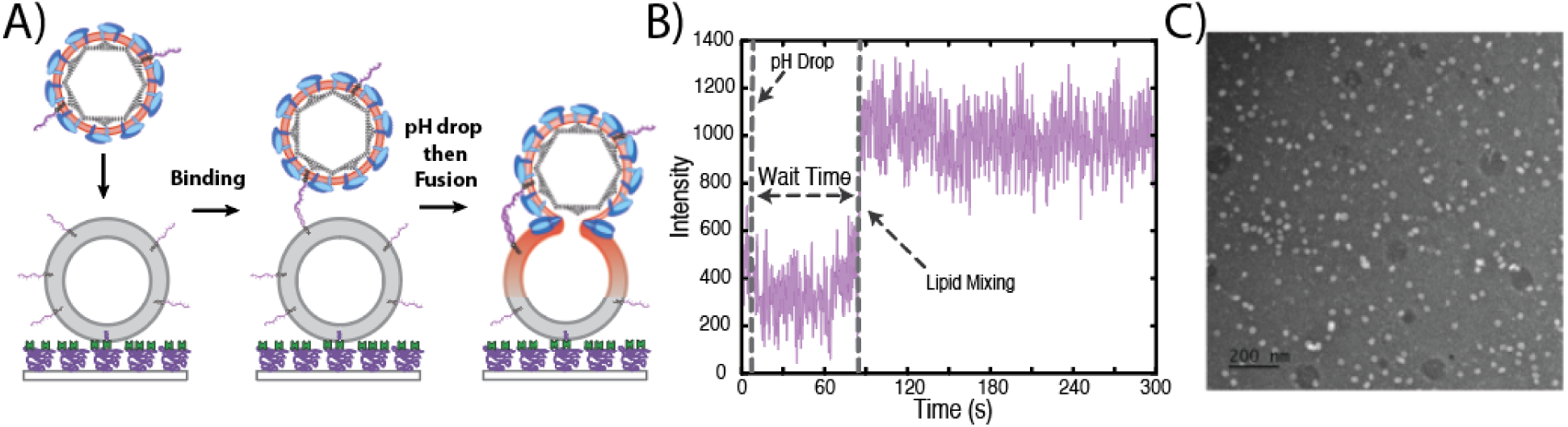
Overview of single VLP fusion assay adapted for DENV2 VLPs. (A) Schematic of single VLP fusion assay. Target liposomes are tethered to a PLL-PEG-coated coverslip inside a PDMS microfluidic channel via Neutravidin-biotin binding. DENV2 VLPs, fluorescently labeled with the Texas Red-DHPE lipid, are bound to the target liposome via DNA-lipid binding in lieu of native receptor binding. Unbound VLPs are removed by rinsing, and then a low pH solution is introduced into the microfluidic channel. VLPs are monitored by fluorescence microscopy, and fusion is identified via fluorescence de-quenching due to lipid mixing between the VLP envelope and target liposome. (B) Example fluorescence intensity trace of a VLP fusing to a target liposome. The fluorescence intensity is calculated as the integrated intensity within a region of interest around the VLP in each video frame. Upon lipid mixing, a sharp spike in the intensity occurs due to fluorescence de-quenching. The time between the pH drop and the onset of lipid mixing is defined as the wait time. (C) Negative stain electron microscopy images of DENV2 VLPs. VLPs are observed to be relatively uniform with diameters ∼20-40 nm. Scale bar is 200 nm.

The distribution of wait times for many VLPs can be visualized in a cumulative distribution function (CDF), such as in **Figure 2a**, and many of the other data figures in this report. The shape and timescale of these distributions are governed by the rate limiting step(s) in the fusion process, and can be used to extract mechanistic information using mathematical modeling (12, 16, 17, 19, 22, 24–26), as described below.

The extent of hemi-fusion is also a useful metric to constrain the mechanistic fusion model, as described below. Extent is measured as the fraction of observed VLPs that underwent fusion during the observation time window. In prior work, we and others have calculated the extent directly from the fusion videos (8, 10, 12, 16, 17). However, only 1 fusion video can be collected per flow cell, which limits the number of VLPs that can be observed. And the light intensity used to capture those few videos is low, introducing additional uncertainty. We therefore developed an improved method which calculates the extent from higher-intensity images taken before and 5 min after the pH drop in several locations around the flow cell which had not otherwise been imaged (see **Supporting Figures S3-S4** for further discussion and validation of this approach). This method was applied to calculate the extents in data throughout this report.

### 2.2. pH-dependence of DENV hemi-fusion kinetics

The pH-dependence of the hemi-fusion kinetics has been essential to propose a robust kinetic model for both ZIKV and WNV (16, 19). To investigate the dependence of dengue virus hemi-fusion on the pH of the triggering buffer solution, we conducted our single VLP fusion assay at pH values from 5.0-6.25 and measured both the distribution of wait times and extents (**Figure 2**). We observed that the wait time distributions (i.e. curve shapes of the CDFs) remained relatively insensitive to pH – all were roughly exponential, with average wait times of ∼30-50 sec (**Figure 2a**). On the other hand, the fusion extents decreased substantially with increasing pH (**Figure 2b**), decreasing by ∼70-80% across the tested pH range. Interestingly, the maximum extent of hemi-fusion (∼0.25 at pH 5) is comparable to that observed for other flaviviruses (∼0.2-0.3 for WNV, ZIKV, see refs (16, 17)).

**Figure 2.**
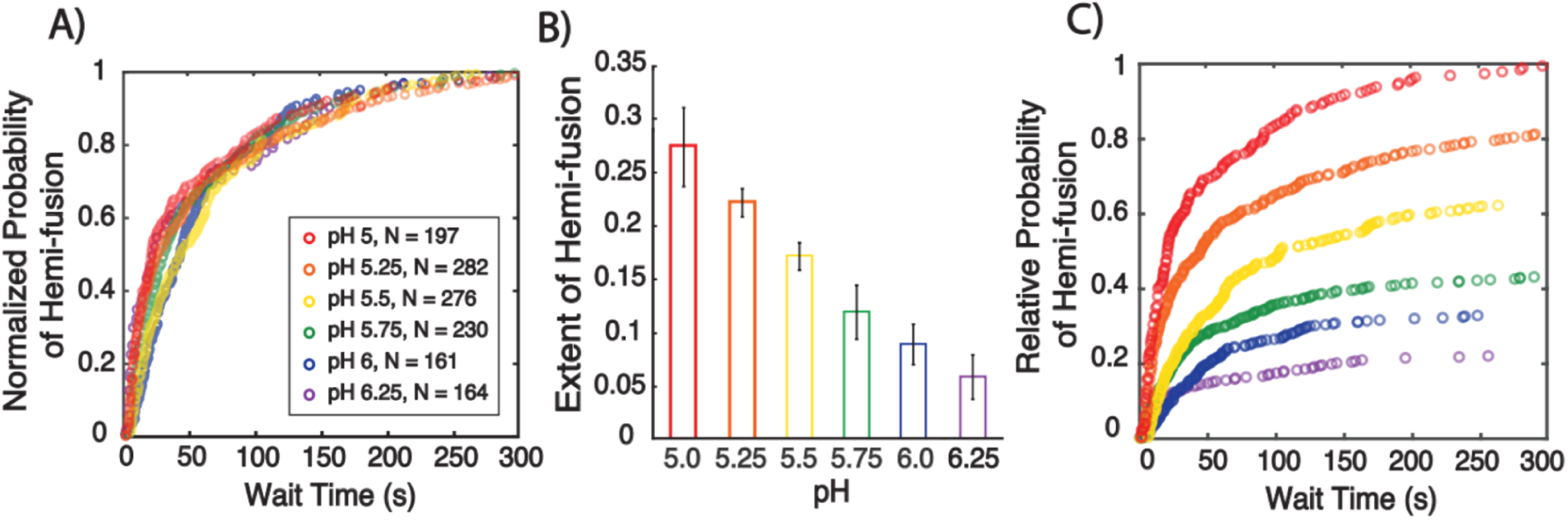
pH dependence of DENV2 VLP hemi-fusion kinetics. Single VLP hemi-fusion experiments were run at differing pH values, ranging from 5 to 6.25. (A) shows the distributions of observed wait times, displayed as cumulative distribution functions (CDFs). CDFs were normalized to the highest value for each data set. (B) shows the extents of hemi-fusion at the different pH values. Extent was measured as the fraction of observed VLPs that underwent hemi-fusion after 5 min following pH drop. Values shown are the mean ± standard deviation of 3 sample replicates. (C) shows the relative distribution of wait times, calculated by multiplying the CDFs in (A) by the mean extents in (B) and normalizing to pH 5.

Together, these data exhibited a strikingly similar pattern to what has been observed previously for single virus measurements of both WNV and ZIKV across similar pH ranges (16, 17, 19): namely, that the hemi-fusion wait time distributions are relatively insensitive to pH (**Figure 2a**), while the extents are strongly sensitive to pH (**Figure 2b**). In the cases of WNV and ZIKV, these observations could not be explained by a traditional linear hemi-fusion model, as demonstrated by mechanistic kinetic modeling (16, 19). Instead, the observations could be explained by a new model which suggested that the rate limiting step of fusion was not pH-dependent across this pH range, and more importantly that the mechanism must contain an off-pathway state that branches from a pH-dependent state.

### 2.3. Kinetic modeling indicates that an off-pathway state exists in the DENV fusion mechanism

To ascertain whether the off-pathway kinetic mechanism developed for WNV and ZIKV (16, 19) could also describe our DENV data, we applied 2 different computational approaches to model the single DENV VLP data. The first approach is “per virus” modeling inspired by chemical kinetics, and the second is “per protein” modeling using a cellular automaton simulation. Each approach is described separately below. In both approaches, we tested the traditional linear model against the off-pathway model.

#### 2.3.1. “Per virus” kinetic modeling supports an off-pathway mechanism

The “per virus” modeling approach is inspired by chemical kinetics and models the entire VLP transitioning between different states leading to hemi-fusion (see reaction schemes in **Figure 3a and b**). This model can in principle accommodate many different states, but the most parsimonious version includes only 3 (linear model) or 4 (off-pathway model).

**State definitions:** All VLPs start in the bound state (State B) at t = 0; this models the VLPs as bound to the target liposomes when the pH drop occurs. Subsequent pH activation allows the VLPs to enter State A, which represents the VLPs in a pH-activated state that can transition to hemi-fusion (State HF) irreversibly. The off-pathway model (**Figure 3b**) also includes State O, which is the off-pathway state. This off-pathway state branches directly from the bound state, putting it in competition with the pH-activation step which leads forward to fusion. This competition between the off-pathway state and pH-activation is required; it leads to the observed decrease in the extent of hemi-fusion as a function of pH.

**Fitting process:** As was done previously for ZIKV and WNV, the *k_AB_* rate constant was tied to the *k_BA_* constant as *k_BA_ × 10^-6.8^*. This enforces a pKa of the activation step of 6.8, consistent with changes in the hydrodynamic radius of Kunjin virus as a function of pH, determined by DLS (17). All other rate constants were fitted as free parameters in the model to the entire data set (wait time distributions and fusion extents for all pH values) simultaneously using maximum likelihood estimation.

**Modeling results:** Consistent with the prior modeling of WNV and ZIKV (16, 19), we observed that the linear model did a very poor job in modeling the DENV data (**Figure 3c-e**). It was able to capture the pH-sensitivity of the fusion extents moderately well (**Figure 3e**), but only at the expense of a very poor quality fit to the wait time distributions (**Figure 3d**). Indeed, this was a consistent problem with the linear model in the WNV and Zika virus reports as well (16, 19). The linear model can only achieve less than maximum extents by decreasing the overall reaction rate such that the process does not level off during the timescale of the measurement. However, this does not match the observed wait time distributions, which clearly show a leveling off in the latter portion of the measurement.

**Figure 3.**
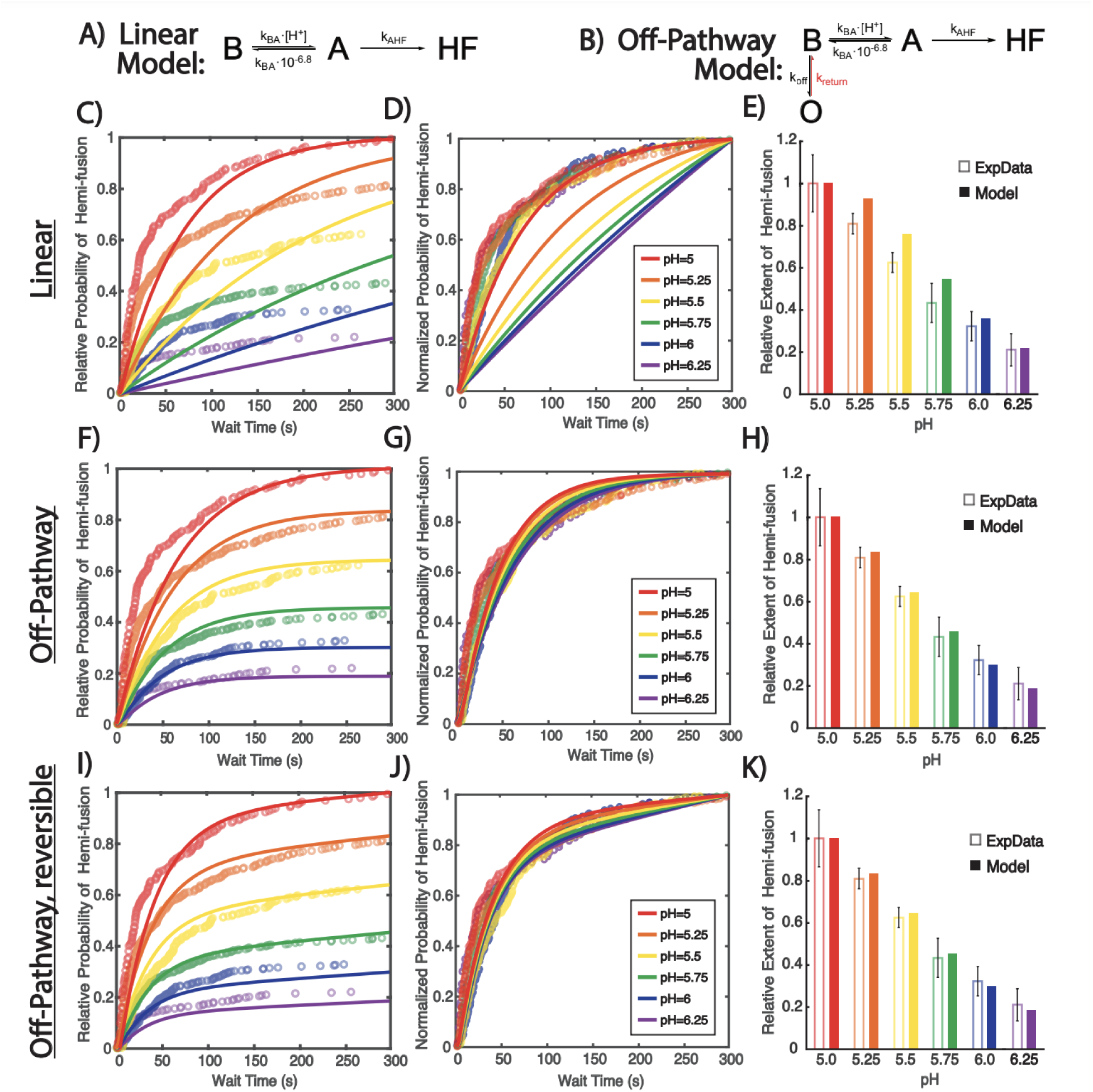
Per virus kinetic modeling supports off-pathway hemi-fusion mechanism. DENV VLP hemi-fusion data collected at different pH values (see Figure 2) was fit to either a linear (A) or an off-pathway (B) mechanism, using “per virus” modeling (state definitions are described in the main text). CDFs of all pH values were fit simultaneously. Panel C shows the best fit (solid lines) to the experimental hemi-fusion CDF (open circles) for the linear model. Panels F and I show the best fits for the irreversible (k_return_=0) and reversible (k_return_≠0) off-pathway models, respectively. The probability of hemi-fusion was calculated relative to the final extent at pH 5, which was set at 1. Panels D, G, and J show the normalized fits on the normalized CDFs to demonstrate how well the fits model the experimental wait time distributions. CDFs were normalized to the final extent of hemi-fusion at each pH value. Panels E, H, and K show the model predictions for final extents of hemi-fusion (solid bars) at t = 300 sec compared to the experimental final extents (open bars). Final extents are shown relative to pH 5, which was set 1. Error bars are the standard deviation of 3 sample replicates. The best-fitting rate constants were: Linear: k_BA_ = 1.4 x 10^3^ M^-1^s^-1^, k_AHF_ = 0.99 s^-1^; Off-pathway: k_BA_ = 8.1 x 10^4^ M^-1^s^-1^, k_off_ = 0.1 s^-1^ and k_AHF_ = 1.4 x 10^-2^ s^-1^; and Off-pathway, reversible: k_BA_ = 6.4 x 10^4^ M^-1^s^-1^, k_off_ = 0.2 s^-1^, k_return_=0.001 s^-1^, and k_AHF_ = 2.4 x 10^-2^ s^-1^.

On the other hand, we observed that the off-pathway model was able to capture the general trends of the DENV data (**Figure 3f-h**). The wait time distributions showed little sensitivity to the pH, matching the experimental data (**Figure 3g**), and the hemi-fusion extents fell within the experimental error for all pH values (**Figure 3h**). By AIC comparison, the relative likelihood of the linear model compared to the off-pathway model approached zero. This supports the conclusion that an off-pathway state is needed to describe the DENV fusion data. Interestingly, all fitted rate constants were within an order of magnitude of the best fitting models reported for both ZIKV and WNV (16, 19).

The off-pathway state need not necessarily be irreversible, as modeled above, although the rate constant of the return (*k_return_* in **Figure 3b**) must necessarily be slow relative to the experimental timescale in order to produce the observed data. We therefore examined whether the off-pathway state might be reversible by incorporating *k_return_* into the model. We observed moderate, but clear improvement to the model fits (**Figure 3i-k**), particularly to the wait time distribution shapes (**Figure 3j**) suggesting that the off-pathway state may be reversible to some extent. AIC analysis supported this conclusion; the relative likelihood of the irreversible off-pathway model was <10^-10^ compared to the reversible model.

#### 2.3.2. “Per protein” kinetic modeling also supports an off-pathway mechanism

The “per protein” approach uses a cellular automaton simulation, which we have previously described (19). Briefly, the simulation models the E proteins of an individual VLP-target liposome interface as a hexagonal lattice. The proteins can transition between the different mechanistic states in the model (see reaction schemes in **Figure 4a and b**), with many states requiring interactions between neighboring proteins. If the right conditions are met during the simulation, the VLP undergoes hemi-fusion and the wait time is recorded. Hundreds of VLPs are simulated at each pH value to model the experimental data.

**State definitions:** The E proteins are initially arranged in dimer pairs in the folded state (State F), from which they can be pH-activated into an extended state (State E). Cooperativity is modeled between the dimer pairs – if one monomer is extended, the rate constant of pH activation for its partner is increased as *CF × k_ac_*_t_., where *CF* is the cooperativity factor. The trimer state (State T) requires 3 neighboring E proteins to be in the extended state (State E), and is also pH-dependent. In order for the VLP to achieve hemi-fusion (State HF), a minimum number of neighboring trimers (*N_tri_*) is required. The off-pathway state (State O) is modeled as branching in competition with the trimerization step; each monomer could enter State O independently. Additional details about the simulation can be found in **Section 4.13** of the Materials and Methods.

**Fitting process:** Several parameters were held fixed during the fitting process, with constraints based on previous data. Analogous to the “per virus” approach, the *k_ret_* rate constant was set at *k_act_ × 10^-6.8^*. *N_tri_* was fixed at 2, the optimal number used for both WNV and ZIKV (16, 19), and In the WNV report, *CF* was observed to not influence the fitting to an appreciable extent, so it was held fixed at 4, the optimal number found for WNV (19). *k_untrimer_* was fixed at 10^-9^ s^-1^, essentially irreversible, consistent with liposome co-flotation data indicating that trimer formation following fusion loop capture is essentially irreversible (17, 27–29). All other parameters were determined by fitting each model to the experimental data using maximum likelihood estimation (see **Section 4.13** in the Materials and Methods).

**Modeling results:** In general, we observed similar results with this modeling approach as we did with the “per virus” approach (compare **Figures 3 and 4**). The linear model produced a very poor fit to the data, unable to capture both the shape of the wait time distribution and the pH-dependent decrease in extent (**Figure 4c-e**). On the other hand, the off-pathway model did, generally speaking, reproduce those features of the data (**Figure 4f-h**). Within the best fitting model, the shape of the wait time distributions were reproduced reasonably well (**Figure 4g**) and the model predictions for the extents showed general agreement with the experimental trend (**Figure 4h**). AIC comparison indicated that the relative likelihood of the linear model compared to the off-pathway model approached zero. When the off-pathway state was treated as reversible, we observed minor, but measurable, improvement to the model (**Figure 4i-k**); relative likelihood of the irreversible vs reversible model was ∼10^-6^ by AIC.

Together, both modeling approaches agree with the conclusions from the previous ZIKV and WNV reports (16, 19); namely, a linear model is insufficient to capture the salient features of the data. Instead, a mechanism that includes an off-pathway state is required. Notably however, the “per protein” approach consistently underperformed relative to the “per virus” approach, suggesting that further model development is needed in the cellular automaton simulations.

**Figure 4.**
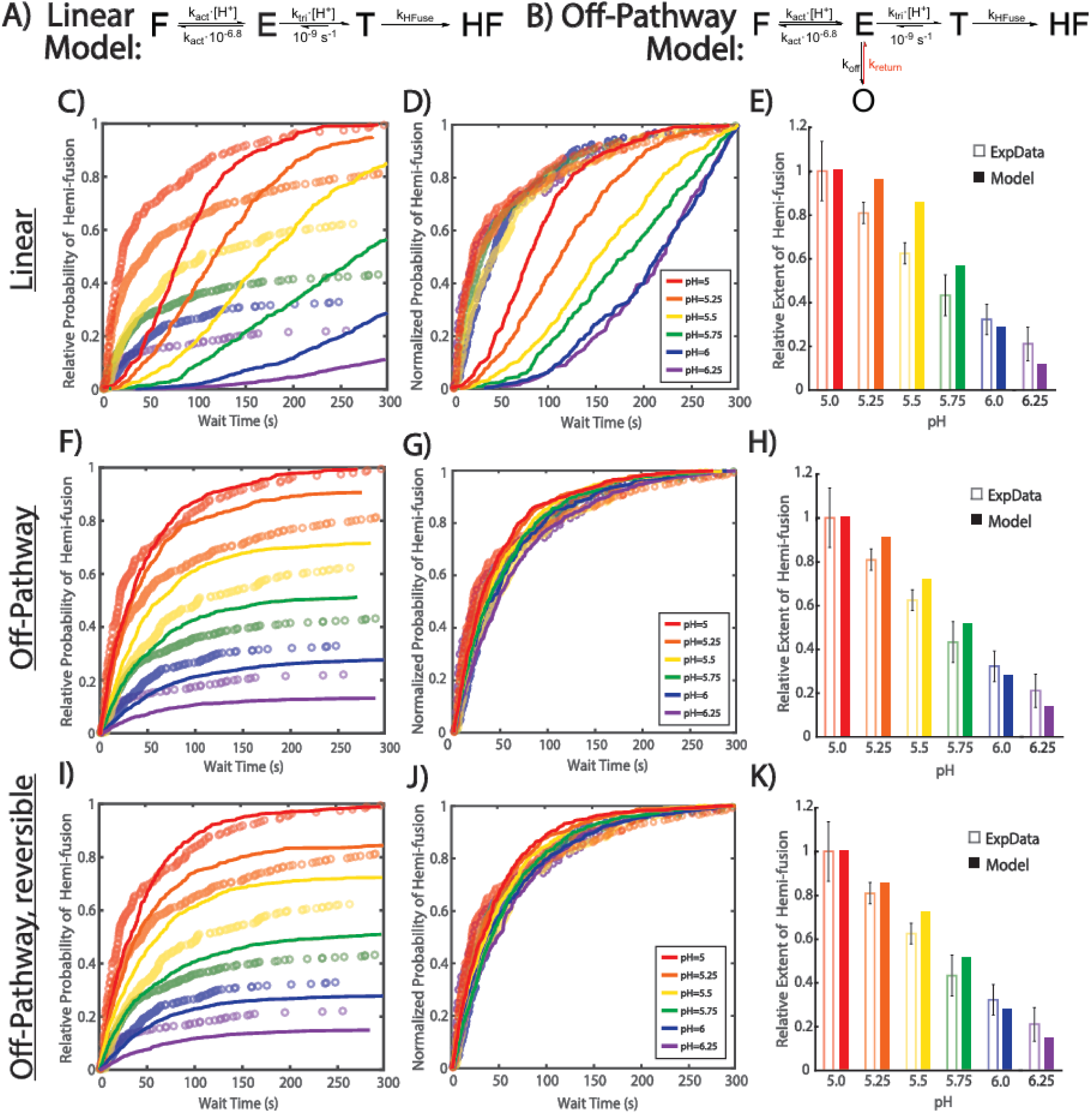
Per protein kinetic modeling also supports off-pathway hemi-fusion mechanism. DENV VLP hemi-fusion data collected at different pH values (see Figure 2) was fit to either a linear (A) or an off-pathway (B) mechanism, using “per protein” cellular automaton simulations (state definitions are described in the main text). CDFs of all pH values were fit simultaneously. Panel C shows the best fit (solid lines) for the linear model to the experimental CDF (open circles). Panels F and I show the best fits for the irreversible (k_return_=0) and reversible (k_return_≠0) off-pathway models, respectively. The probability of hemi-fusion was calculated relative to the final extent at pH 5, which was set at 1. Panels D, G, and J show the normalized fits on the normalized CDFs to demonstrate how well the fits model the experimental wait time distributions. CDFs were normalized to the final extent of hemi-fusion at each pH value. Panels E, H, and K show the model predictions for final extents of hemi-fusion (solid bars) at t = 300 sec compared to the experimental final extents (open bars). Final extents are shown relative to pH 5, which was set 1. Error bars are the standard deviation of 3 sample replicates.The best-fitting rate constants were: Linear: k_act_ = 1.0 x 10^5^ M^-1^s^-1^, k_tri_ = 1.0 x 10^3^ M^-1^s^-1^, k_hemi_ = 7.0 x 10^-3^ s^-1^ Off-pathway: k_act_ = 8.0 x 10^8^ M^-1^s^-1^, k_tri_ = 8.0 x 10^6^ M^-1^s^-1^, k_off_ = 25.3 s^-1^, k_hemi_ = 7.5 x 10^-3^ s^-1^ and Off-pathway, reversible: k_act_ = 8.0 x 10^8^ M^-1^s^-1^, k_tri_ = 8.0 x 10^6^ M^-1^s^-1^, k_off_ = 25.3 s^-1^, k_hemi_ = 7.5 x 10^-3^ s^-1^, and k_return_=0.12 s^-1^.

### 2.4. Reversibility of the off-pathway state

In both our modeling approaches described above, we observed moderate to substantial improvement when the off-pathway state was modeled as reversible. To explore this experimentally, we measured the extent of hemi-fusion across a longer time window, up to 120 minutes following pH drop at pH 5. We observed a slow increase in the extent of hemi-fusion up to ∼20 minutes, where it leveled off at ∼0.3 (**Figure 5**).

We then compared the predictions of our best fitting “per virion” model (i.e. **Figure 3i-k** above) to this long timescale data, varying the reversibility rate constant, *k_return_* (**Figure 5**). As expected, we observed poor agreement with the data when the off-pathway state was modeled as irreversible. On the other hand, the reversibility rate constant from our prior best fit (*k_return_* = 0.001 s^-1^) only modeled the data well up until ∼20 min. A better match was observed when the rate constant was increased to 0.004 s^-1^. This data (as well as both our modeling approaches above) indicate that there is slow reversibility from the off-pathway state, allowing at least some VLPs to progress to hemi-fusion over a longer time window.

Finally, it is also notable that the extent values in the long timescale data level off well below extent = 1 (**Figure 5**), suggesting that a large fraction of the VLPs may be fusion-incompetent to begin with, perhaps due to differing levels of maturation during production or subsequent inactivation during the isolation process. Alternatively, although not mutually exclusive, some VLPs may get trapped in a more irreversible off-pathway configuration which is not yet captured by the simplistic models.

**Figure 5.**
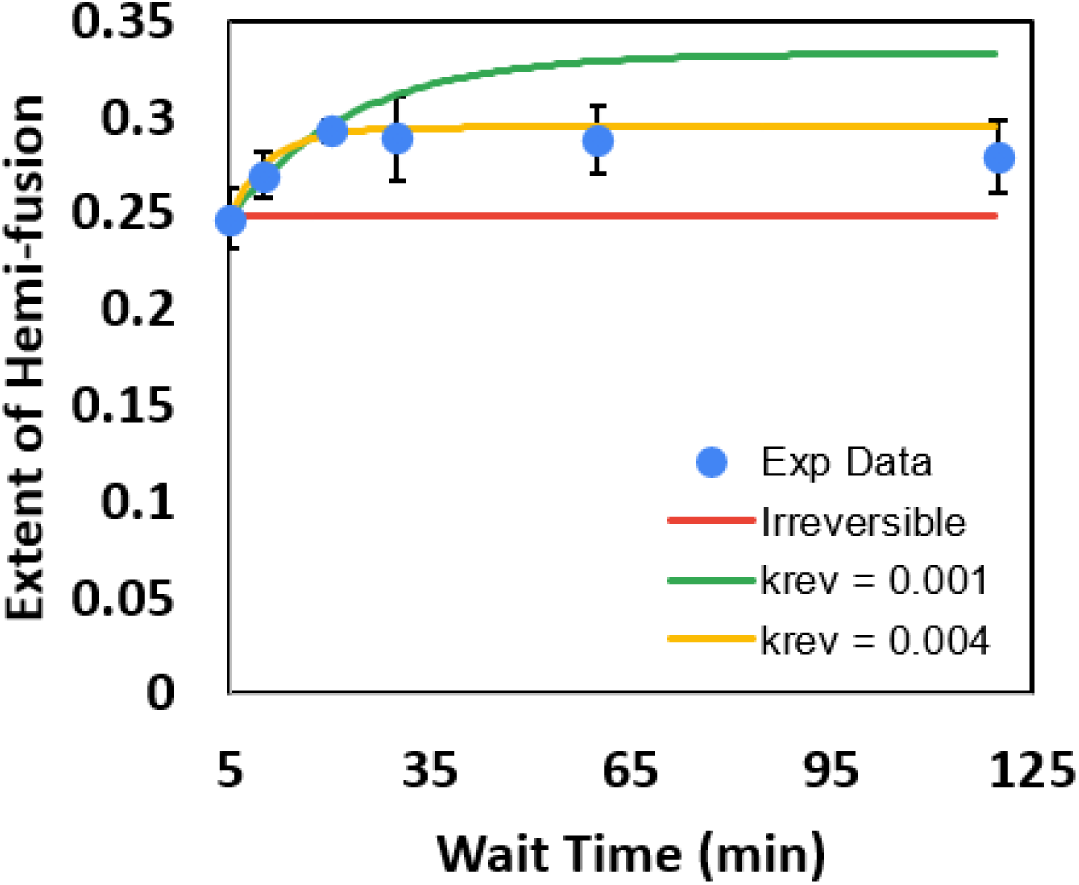
Reversibility of the off-pathway state is supported by long timescale hemi-fusion extents. Single VLP hemi-fusion was monitored by time-lapse microscopy at pH 5 up to 120 minutes following pH drop. The measured hemi-fusion extents at each time point are shown (solid dots, representing mean ± standard deviation of 3 different image regions within the sample). This data was compared to predictions of the best fitting “per virus” model (see Figure 3i-k) with k_return_ fixed at 0 s^-1^ (red line, irreversible model), 0.001 s^-1^ (green line), or 0.004 s^-1^ (gold line). Model predictions were calculated relative to the experimental extent at 5 min, consistent with the prior analysis in Figure 3 above. The other rate constants employed were k_BA_ = 6.4 x 10^4^ M^-1^s^-1^, k_off_ = 0.2 s^-1^, and k_AHF_ = 2.4 x 10^-2^ s^-1^.

### 2.5. Influence of late endosomal (LE) lipid composition on DENV hemi-fusion

In the hemi-fusion data described above (e.g. **Figures 2, 5**), the target liposomes were composed of a lipid mixture that mimics the late endosomal (LE) membrane (48.95% POPC, 20% DOPE, 20% BMP, 10% PI, 1% biotinyl PE, 0.05% Oregon Green-DHPE, see ref (30)), which is the physiological site of DENV fusion (31, 32). Previous work has indicated that specific late endosomal lipid components, especially anionic lipids such as BMP, may be important to ensure that DENV fusion occurs in this cellular compartment (30). However, in previous single virus fusion studies of other flaviviruses, lipid mixtures were employed that did not include late endosomal anionic lipids: the ZIKV study cited above (16) used 69.9% POPC, 20% DOPE, 10% Chol, 0.01% Oregon Green-DHPE, while previous WNV and DENV studies (17, 18) have used 40% POPE, 18.2% POPC, 18.2% DOPC, 18.2% Chol, 0.2% Ni-NTA-DOGS, 9.1% carboxyfluorescein-PE. To test whether the LE composition would influence the fusion behavior in our assay, and in particular whether evidence of an off-pathway state would still be observed, we collected a smaller single VLP fusion data set (pH 5 and 5.5 only) using an adaptation of the composition used previous for ZIKV (simply swapping BMP and PI for cholesterol).

Interestingly, we observed nearly identical results as with the LE mixture. The wait time distributions at pH 5 and 5.5 overlapped quite well, and showed nearly identical curve shapes with data collected using the LE mixture (**Figure 6a**). Similarly, the hemi-fusion extents were within error when comparing the 2 mixtures (**Figure 6b**).

Together, this data indicates that in our assay the rate-limiting step of hemi-fusion as well as the requirement for an off-pathway state in the mechanism are not dependent on the late endosomal lipid mixture, including BMP. It is possible that anionic lipids such as BMP may be more important for the transition from hemi-fusion to pore opening (full fusion) in DENV fusion, which is not detected in our lipid mixing assay. This interpretation would be consistent with the conclusions of prior work of DENV fusion to liposomes in bulk (30).

**Figure 6.**
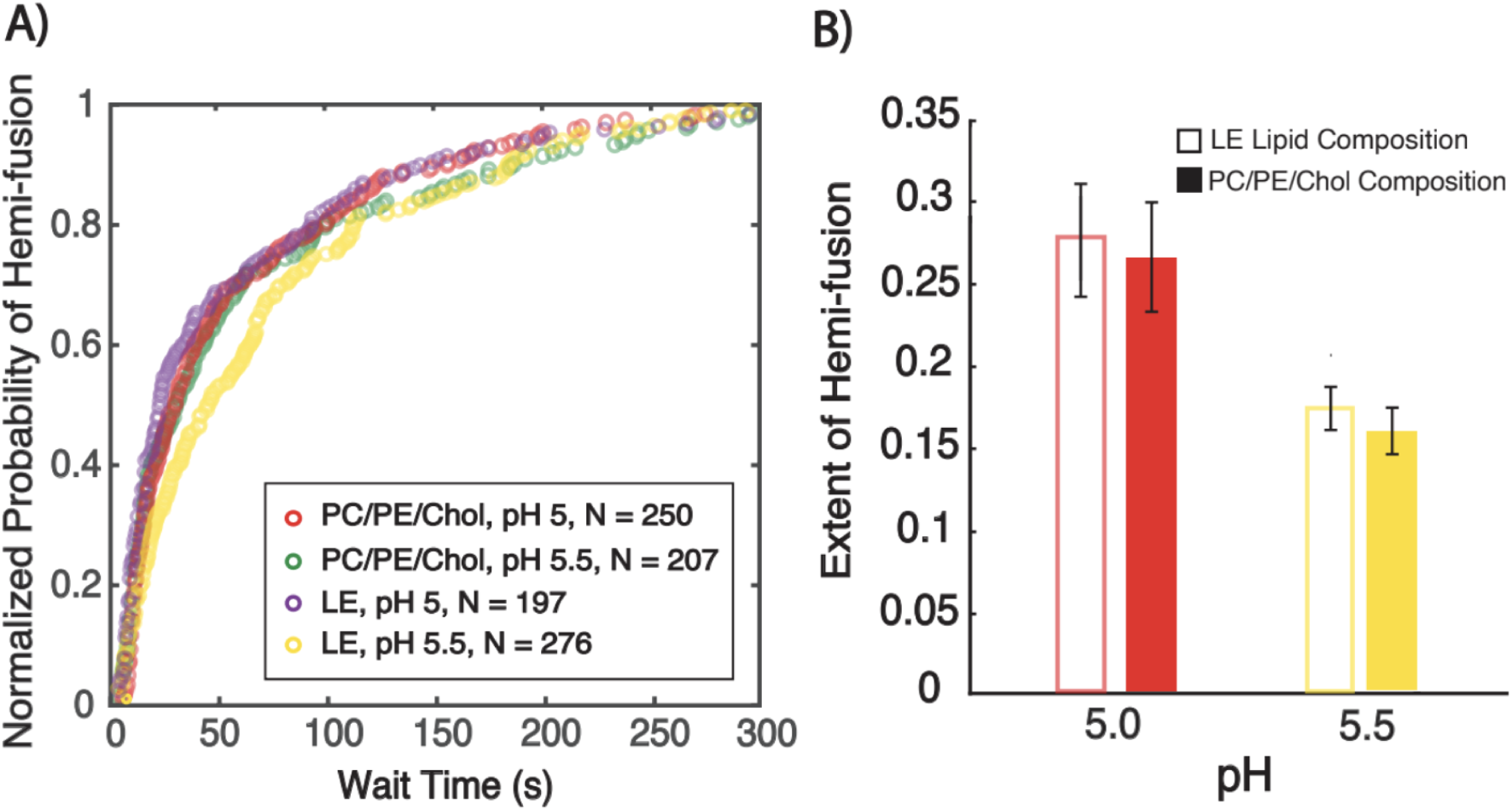
Influence of late endosomal lipid components on DENV2 hemi-fusion. Single VLP hemi-fusion experiments were run at pH 5 and 5.5 using target liposomes with a late endosome (LE) mimic composition or a PC/PE/Chol lipid mixture, in which the BMP and PI lipids had been swapped with cholesterol. (A) shows the distributions of observed wait times, displayed as cumulative distribution functions (CDFs). CDFs were normalized to the highest value for each data set. (B) shows the extents of hemi-fusion at the different pH values. Extent was measured as the fraction of observed VLPs that underwent hemi-fusion after 5 min following pH drop. Values shown are the mean ± standard deviation of 3 sample replicates. LE composition = 48.95% POPC, 20% DOPE, 20% BMP, 10% PI, 1% biotinyl PE, 0.05% Oregon Green-DHPE. PC/PE/Chol composition = 48.95% POPC, 20% DOPE, 30% cholesterol, 1% biotinyl PE, 0.05% Oregon Green-DHPE.

## 3. Conclusion

We have utilized single VLP lipid mixing measurements to study the pH dependence of the kinetics of DENV hemi-fusion. This data, coupled with our computational modeling, provides strong evidence of an off-pathway state in the mechanism of DENV hemi-fusion. When considered together with prior work (16, 19) demonstrating that both ZIKV and WNV also follow an off-pathway fusion mechanism, this indicates that an off-pathway state may be a feature of flavivirus fusion more broadly. Additionally, the fitted rate constants for the best fitting models of all three viruses are all very similar (approximately within an order of magnitude of each other), underscoring the similarities in the fusion mechanism employed by these viruses.

This report also provides some additional nuance to this off-pathway state, at least for DENV. First, by using long timescale measurements of hemi-fusion extent, we have identified that the off-pathway state appears to be reversible over the timescale of tens of minutes, at least for some fraction of virions. Second, we have provided evidence that late endosomal anionic lipid components such as BMP in the target membrane do not appear to be required for this off-pathway mechanism, as their presence or absence did not substantially alter the observed hemi-fusion data.

The precise nature of this off-pathway state cannot be determined from the data in this report. In prior work, we have postulated that this off-pathway state may represent misfolded or aggregated E monomers either before or after membrane insertion, or a fusion incompetent trimer state following the dimer-to-trimer transition. Disentangling these structural hypotheses will be the focus of future work, and will likely require both new single VLP fusion data, computational modeling, and additional biochemical data.| Furthermore, small molecule inhibitors of flavivirus fusion developed by our group and others may be useful in probing the fusion pathway. We also note that there is the potential to use pharmacological approaches to stabilize the off-pathway state as an antiviral strategy.

## 4. Materials and Methods

### 4.1. Materials

Dioleoylphosphatidylethanolamine (DOPE), 1,2-dioleoyl-sn-glycero-3-phosphocholine (DOPC), 1,2-dipalmitoyl-sn-glycero-3-phosphoethanolamine-N-biotinyl (biotinyl PE), Bis(Monoacyl-glycero)Phosphate (BMP), cholesterol (chol) and 1-stearoyl-2-hydroxy-sn-glycero-3-phosphoinositol (LysoPI) were purchased from Avanti Polar Lipids (Alabaster, AL). Poly-L-Lysine graft PEG polymer (PLL(20)-g[3.5]-PEG(2), abbreviated PLL-g-PEG) and Poly-L-lysine graft biotinylated PEG polymer (PLL(20)-g[3.5]-PEG(2)/PEG(3.4)-biotin(50%), abbreviated PLL-g-PEG-biotin) were purchased from SuSoS AG (Dübendorf, Switzerland). Pluronic™ F-127 (20% Solution in DMSO), Neutravidin protein, Oregon Green-1,2-dihexadecanoyl-sn-glycero-3-phosphoethanolamine (OG-DHPE) and Texas Red-1,2-dihexadecanoyl-sn-glycero-3-phosphoethanol-amine (TR-DHPE) were obtained from Thermo Fisher Scientific (Waltham, MA, USA). Polydimethylsiloxane (PDMS) Sylgard 184 elastomer base and curing agent were purchased from Ellsworth Adhesives (Germantown, WI, USA). Chloroform, methanol, and buffer salts were obtained from Fisher Scientific (Pittsburgh, PA) and Sigma-Aldrich (St. Louis, MO, USA). DNA-lipids were a gift from Prof. Steven Boxer at Stanford University, prepared by phosphoramidite coupling of short DNA oligos to a double-tailed C18 lipid as previously described (20, 33). DNA-lipid sequences used were (listed 5’ to 3’ direction) 5’A24 = lipid - AAA AAA AAA AAA AAA AAA AAA AAA; 5’T24 = lipid - TTT TTT TTT TTT TTT TTT TTT TTT. DNA-lipid concentration was determined by UV-visible spectroscopy using the calculated molar extinction coefficients of the DNA sequence at 260 nm (the lipid tail absorbs negligibly at this wavelength).

### 4.2. Buffer Definitions

All buffers were filtered at 0.22 μm.

- Reaction buffer (RB pH 7.4) = 10 mM NaH2PO4, 90 mM sodium citrate, 150 mM NaCl, pH 7.4.
- TrN buffer (TrN) = 20 mM Tricine-HCl, 140 nM NaCl, 0.005% Pluronic F-127, pH 7.8.
- Sodium Acetate buffer (SA, pH 5.0 to 5.50 as indicated) = 140 mM NaCl, 20 mM Tricine-HCl, 100 mM sodium acetate, 0.005% Pluronic F-127, titrated with HCl and/or NaOH to the indicated pH.
- MES buffer (MES, pH 5.75 to 6.25, as indicated) = 140 mM NaCl, 20 mM Tricine-HCl, 100 mM MES, 0.005% Pluronic F-127, titrated with HCl and/or NaOH to the indicated pH.
- HEPES Buffer (HB) = 20 mM HEPES, 150 mM NaCl, pH 7.2

### 4.3. Fluorescence Microscopy

A Zeiss Axio Observer 7 microscope (Carl Zeiss Microscopy, LLC., White Plains, NY) was used to acquire fluorescence microscopy images. The microscope was equipped with a 63x oil immersion objective, NA=1.4, a Lumencor Spectra III, LED Light Engine as an excitation light source, and a Definite Focus 3. Images were recorded with a Hamamatsu ORCA Flash 4.0 V2 Digital CMOS camera (Hamamatsu Photonics K.K., Hamamatsu City, Japan) with a 16-bit image setting. The microscope was operated using Micromanager software (34). Images and video micrographs were captured at 100 or 300 ms/frame with 2x2 binning. Texas Red images were obtained using the following filter cube: ex = 562/40 nm, bs = 593 nm, em = 641/75 nm, light engine typical intensity setting = 10/1000 or 25/1000 (green LED). Oregon Green images were captured using the following excitation/emission filter cube: ex = 475/50 nm, bs = 506 nm, em = 540/50 nm, light engine typical intensity setting = 2/1000 (cyan LED)

### 4.4. Preparation of Dengue Virus Like Particles (DENV VLPs)

Hek293 cells were cultured in Dulbecco’s Modified Eagle Medium (DMEM) supplemented with 10% FBS, 10 mM HEPES, and 1X MEM Non-Essential Amino Acids. To produce VLPs, the cells were seeded to 80% final confluence in DMEM + 2% FBS then transfected with the pcDNA3.1 vector with a structural cassette containing the codon-optimized sequence of DV2-FGA/02 prM-E described by Wang and colleagues using Lipofectamine 3000 (35, 36). The transfected cells were incubated for 48 h at 28°C before the supernatants were collected. Cells were then removed by centrifugation and the supernatant precipitated while continuously rotating for 3 h at 4°C by addition of 8% (wt/vol) of polyethylene glycol 8000 (PEG 8000) and 0.5M NaCl.. The VLPs were pelleted by centrifugation at 10,000 × g for 30 min at 4°C. The pellet was resuspended in HB buffer and processed for electron microscopy. The VLP samples were attached to glow-discharged 300 mesh carbon/formvar coated Cu grids and stained with 1% Uranyl Acetate in H_2_O using single-droplet negative staining. Samples were imaged using a JEM-1400 electron microscope (Jeol USA, Peabody, MA).

### 4.5. Fluorescence labeling of DENV VLPs and DNA-lipid incorporation

To prepare the dye labeling solution, 22 µL of 0.75 g/L TR-DHPE dye in ethanol was combined with 0.5 µL Pluronic F-127 (20% solution in DMSO), vortexed thoroughly and sonicated at RT for 5 minutes. 2 µL of dye labeling solution was added to 50 µL of VLPs (typically at ∼0.9 g/L total protein determined by BCA assay), with a final TR-DHPE concentration of 21 µM. This solution was incubated at room temperature (RT) for 30 minutes, and then at 4 ℃ for two hours. Labeled VLPs were purified from free dye by size exclusion gravity chromatography, using G-100 sephadex beads equilibrated in TrN buffer. Fractions containing labeled VLPs were stored in TrN buffer at -80°C until used. Note that in limited data sets we found that long-term storage in a slightly basic buffer (pH 7.8 in this case) may be important to not inactivate the VLPs, which presumably occurs due to low levels of premature triggering of the viral E protein at neutral pH. This was not tested exhaustively however. Prior to use in fusion assays, labeled VLPs were thawed and syringe-filtered through a Millex-GV 0.22 um PVDF low binding filter (MilliporeSigma, Burlington, MA, USA). The approximate VLP concentration was determined by “splat” assay (described below). 5’A24 DNA-lipid was then added at a ratio of 10 DNA-lipids per VLP (see Figure SX for how this ratio was determined), and the mixture was incubated at RT for 30 minutes to incorporate into the VLP membrane. Previous work has demonstrated that such incubation produces quantitative transfer of the DNA-lipids into the viral membranes (10, 20, 23).

### 4.6. Microfluidic flow cell preparation

Microfluidic flow cells were constructed of polydimethylsiloxane (PDMS) chips plasma bonded to clean glass coverslips, as previously described (20). Briefly, PDMS chips were constructed with 2 parallel flow channels (each was 2.5 mm x 13 mm x 70 μm) by pouring a 10:1 mixture of Sylgard 184 elastomer base and curing agent onto an appropriate mold made by “tape-based soft lithography” (37), and then curing for at least 2 hours at 70°C. Individual chips were cut out with a scalpel, and inlet/outlet holes were created using a biopsy hole puncher (2.5 mm diameter, Harris Uni-core, Ted Pella Inc.). Separately, glass coverslips (24 x 40 mm, NO. 1.5 VWR International, Randor, PA) were cleaned by first immersing in a 1:7 solution of 7x detergent (MP Biomedicals, Burlingame, CA) and deionized water, and heating to clarity. The coverslips were then rinsed extensively under deionized water, followed by Milli-Q water for two minutes, and then baked in a kiln at 400 °C for 4 hours. To assemble the flow cell, PDMS chips and cleaned coverslips were activated by exposure to air plasma for 70 seconds in a Harrick Plasma Cleaner PDC-3xG (Harrick Plasma, Ithaca, NY). After plasma exposure, the PDMS chip was immediately pressed, channels facing down, onto the cleaned glass coverslip to create a plasma bond between the two. A buffer well cut from the end of a 1 mL plastic pipette tip was then affixed to the inlet hole of each channel with five-minute epoxy (Devcon, ITW Polymer Adhesives North America, Danvers, MA).

### 4.7. “Splat” assay to determine approximate VLP concentration

The approximate concentration of labeled VLPs was determined using a “splat” assay, which we have described previously for use with fluorescently labeled virions (38). Briefly, Texas Red-labeled VLPs were mixed 1:1 with Oregon Green-labeled liposomes at a low concentration (typically 1:2000 dilution of liposomes at 0.4 mg/mL). This mixture was then added to a freshly prepared microfluidic flow cell, and incubated for several minutes at RT. Under such conditions, both the VLPs and liposomes quickly adhere (or “splat”) to the glass coverslip. Fluorescence micrographs of the VLPs and liposomes were collected in separate fluorescence channels, and the numbers of particles in each image were quantified using custom-written Matlab scripts (source code available at https://github.com/rawlelab/Flavi-Analysis-Modeling-Public). The nominal concentration of the liposomes can be determined by straightforward back of the envelope calculations (38). Then, the approximate VLP concentration can be calculated simply as the ratio of observed VLPs to liposomes in a field of view, assuming that the VLPs and liposomes both bind to the glass coverslip quantitatively. Using this method, typical concentrations of labeled VLP fractions were found to be ∼tens of pM, after accounting for dilutions during preparation.

### 4.8. Target liposome preparation

Liposomes were prepared by thin-film hydration followed by extrusion (39). Briefly, a lipid mixture was prepared in chloroform/methanol with the desired molar ratio of lipid components (1.4 x 10^-7^ moles of total lipid). Unless noted otherwise, the standard composition for tethered liposomes was 48.95 mol% DOPC, 20 mol% DOPE, 20 mol% BMP, 10 mol% LysoPI, 1 mol% biotinyl PE, and 0.05 mol% OG-DHPE. The organic solvent was evaporated under a stream of N_2_ gas, and the resulting lipid film was then dried under house vacuum for at least two hours. The dried film was hydrated in 250 μL reaction buffer (RB pH 7.4) for ten minutes and then resuspended by vortexing at maximum speed for at least sixty seconds. This suspension was extruded at least 21 times using a mini-extruder (Avanti Polar Lipids, Alabaster, AL) with a 100 nm-pore membrane to produce large unilamellar liposomes. The average diameter of liposomes was measured to be ∼120-130 nm by dynamic light scattering (DynaPro NanoStar DLS, Wyatt Technologies, Santa Barbara, CA). Liposomes were stored at 4°C and used within approximately a week.

DNA-lipids were incorporated into liposomes to mediate binding with virus-like-particles (see schematic in **Figure 1a**). 5’T24 DNA-lipid was added at 0.05 mol % (or ∼65 DNA/liposome for a 120 nm liposome) to the desired volume of liposomes and stored at 4°C overnight to allow for complete insertion into the liposome outer leaflet.

### 4.9. Tethering liposomes inside microfluidic device

Liposomes were tethered inside a microfluidic device by biotin-NeutrAvidin binding between biotinylated lipids in the liposome and a biotinylated polymer affixed to the glass coverslip (see schematic in **Figure 1a**). To prepare the polymer surface, 5 μL of a 95/5 PLL-PEG/PLL-PEG-biotin solution was added to each channel of the microfluidic device and incubated for 30 minutes at room temperature. Each channel was rinsed with 1.5 mL of milliQ water followed by 1.5 mL of RB (pH 7.4) using a Fusion 200 syringe pump (Chemyx Inc., Stafford, TX, USA), flow rate 800 µL/min. Next, 6 μL of 0.2 mg/mL NeutrAvidin protein dissolved in RB (pH 7.4) was exchanged into each channel using a pipette, incubated for 15 minutes, and then rinsed with 1.5 mL of TrN (pH 7.8) by syringe pump. Lastly, 6 µL of target liposomes with biotinyl PE (0.4 mg/mL total lipids) was added, incubated for 60 minutes, and then rinsed with 1.5 mL of TrN (pH 7.8) by syringe pump at 800 µL/min to remove any unbound liposomes.

### 4.10. Single VLP Hemi-fusion Assay and Analysis

After visualizing the tethered liposomes by fluorescence microscopy, 4 µL of fluorescently labeled VLPs (typical concentration ∼50 pM) containing DNA-lipids were added to the flow cell and incubated for 10 minutes at room temperature to facilitate binding. Unbound VLPs were removed by rinsing with 1 mL TrN buffer by syringe pump, flow rate 800 µL/min. Proper binding was verified by fluorescence microscopy imaging in the Texas Red channel, and a suitable location was chosen. A hemi-fusion video micrograph was then initiated. Within the first few seconds of the video, the pH was lowered by buffer exchange into the flow cell (SA or MES buffer, pH as indicated), and individual VLPs were then monitored for 5 minutes. Calibration measurements observing the decrease in fluorescence intensity of pH-sensitive Oregon Green-DHPE in supported lipid bilayers demonstrated that the uncertainty in the timing of pH drop using this approach was ∼1-2 seconds, much shorter than the average timescale of hemi-fusion (tens of seconds).

Hemi-fusion video micrographs were analyzed using an adaptation of previously published custom-written Matlab scripts (source code available at https://github.com/rawlelab/Flavi-Analysis-Modeling-Public). These scripts automatically find labeled VLPs, measure their fluorescence intensity over time, and identify sudden jumps in fluorescence indicative of dye dequenching due to lipid mixing. Correct classification of hemi-fusion events and the identification of the video frame in which fusion occurred was verified by the user.

### 4.11. Measuring Hemi-fusion extent using “Before and After” Method

Images of VLPs in several locations throughout the flow cell were collected before the pH drop using 2.5 fold higher light intensity (int = 25/1000 on the Spectra III LED light engine). Subsequently, the pH drop was initiated by buffer exchange, and images in the same locations were again collected after 5 minutes (or longer times in some data sets as indicated). These areas were not exposed to excitation light in the intervening time, although in some instances a hemi-fusion video micrograph was collected in a distant location in the flow cell.

This data was analyzed using custom-written Matlab scripts (source code available at https://github.com/rawlelab/Flavi-Analysis-Modeling-Public). These scripts quantify the integrated intensity of each particle in the before and after images. If the particle’s intensity has increased by a fraction greater than the threshold cutoff (value = 0.4), hemi-fusion is determined to have occurred. This threshold cutoff value was established by collecting and analyzing before/after data of labeled VLPs adhered non-specifically to a clean glass coverslip. A value of 0.4 was selected as the threshold which would produce <2% error in this control sample (see Figure S3).

### 4.12. Per Virus Kinetics Modeling

Per virus kinetic modeling inspired by chemical kinetics was performed using Matlab scripts as previously described to model WNV and ZIKV data (16, 19) (source code available at https://github.com/rawlelab/Flavi-Analysis-Modeling-Public). Briefly, the system of coupled ordinary differential equations representing each kinetic model was solved numerically for a given set of rate constants and evaluated at discrete time points to determine the number of VLPs in each state. At t = 0 seconds, all VLPs were defined to be in state B (bound state). At each subsequent time point, the number of VLPs that had achieved the hemi-fusion state (state HF) was recorded, and compiled into a cumulative distribution function (CDF). An constrained optimization method was used to determine the best fitting rate constants by maximum likelihood estimation as previously described (19). Optimization constraints were: Linear model: *k_BA_* = 1 x 10^2^ to 1 x 10^5^ M^-1^s^-1^, *k_AHF_* = 0.01 to 5 s^-1^; Off-pathway: *k_BA_ =* 1 x 10^4^ to 2 x 10^5^ M^-1^s^-1^, *k_off_* = 0.05 to 0.5 s^-1^ and *k_AHF_* = 0.01 to 0.03 s^-1^; and Off-pathway, reversible: *k_BA_* = 1 x 10^4^ to 2 x 10^5^ M^-1^s^-1^, *k_off_* = 0.05 to 0.5 s^-1^, *k_return_*=1 x 10^-5^ to 0.5 s^-1^, and *k_AHF_* =0.01 to 0.03 s^-1^. CDFs from all pH values were fit simultaneously. During the fitting, *k_AB_* was fixed at *k_BA_ × 10^-6.8^*, as described in the Results and Discussion; all other rate constants were determined as above. In all cases, rate constants converged to values well within the boundary constraints.

### 4.13. Per Protein Kinetics Modeling: Cellular Automaton Simulations

Cellular automaton modeling of single VLP fusion data was performed in Matlab as we have previously described (16, 19) (source code available at https://github.com/rawlelab/Flavi-Analysis-Modeling-Public). Briefly, the E proteins at an individual VLP-target membrane interface were modeled as an hexagonal lattice consisting of 30 monomers, arranged into dimer pairs. Each VLP was modeled for 300 seconds, or until it reached hemi-fusion (State HF). During the simulation, monomers could transition between states in the model only when certain conditions were met, often involving the state of neighboring monomers. Conditions included: 1) all monomers started in the natively folded state (State F) at *t* = 0 s, 2) if one monomer entered the pH-activated extended state (State E), its dimer pair received a “cooperativity boost” modeled as *CF × k_ac_*_t_, 3) three adjacent monomers in the extended state were required to access the trimer state (State T), after which the three monomers functioned as a trimer unit, 4) two adjacent trimers were required to access the hemi-fused state (State HF).

For each pH value and set of rate constants, simulations were run in batches of 50 VLPs until at least 500 VLPs had undergone hemi-fusion, or until 2500 VLPs had been simulated. The minimum simulation time step was 100 ms, although the time step could be larger – the time step was varied with each pH value and set of rate constants such that the maximum transition probability for any transition between states would be 0.1.

Some model parameters (*k_ret_*, *N_tri_*, *CF,* and *k_untrimer_*) were held fixed as described in the Results and Discussion. The remaining parameters were determined by best fits to the experimental data using a grid-based search algorithm and maximum likelihood estimation as previously described (19). Ranges scanned for each parameter were: Linear model: *k_act_*= 1 x 10^4^ to 1 x 10^8^ M^-1^s^-1^ (100x faster than *k_tri_*), *k_tri_* = 1 x 10^2^ to 1 x 10^6^ M^-1^s^-1^, *k_hemi_* = 1 x 10^-4^ to 7 x 10^-2^ s^-1^; Off-pathway: *k_act_* = 1 x 10^7^ to 1.2 x 10^9^ M^-1^s^-1^ (100x faster than *k_tri_*), *k_tri_* = 1 x 10^5^ to 1.2 x 10^7^ M^-1^s^-1^, *pk_off_ =* 5 to 7.5, *k_hemi_* = 1 x 10^-4^ to 7 x 10^-2^ s^-1^; Note that *k_off_* was scanned in log space as *k_off_ = k_tri_ × 10^-pKoff^,* as described previously (19). The reversible off-pathway model was examined using the best fit parameters for the irreversible model and additionally scanning *k_return_* = .025 to 0.25 s^-1^.

## Acknowledgements

The authors thank Prof. Steve Boxer (Stanford University) for the DNA-lipids and Prof. Stephen Harrison (Harvard University) for the DENV VLP plasmid used in this report. RJR acknowledges financial support from Williams College and NIH grant R15AI171754. PLY acknowledges support from NIH R01AI14615206. The project described was supported, in part, by NIH S10 Award Number 1S10OD028536-01, titled “OneView 4kX4k sCMOS camera for transmission electron microscopy applications” from the Office of Research Infrastructure Programs (ORIP). Its contents are solely the responsibility of the authors and do not necessarily represent the official views of the NCRR or the National Institutes of Health.

## Author Contributions

TA, FC, MK designed experiments, collected and analyzed data. CC, BL, and PLY contributed resources. TA also helped write the manuscript. RJR designed experiments, analyzed data, acquired project funding, and wrote the manuscript. PLY acquired funding and provided supervision. All authors reviewed the manuscript.

## Declaration of interests

The authors declare no competing interests.

## Notes

### Competing Interest Statement

The authors have declared no competing interest.

