## Supplemental Information for "Single VLP lipid-mixing measurements confirm off-pathway state in dengue virus fusion mechanism"

**Affiliations:**
[1] Department of Chemistry, Williams College, Williamstown, MA
[2] Department of Microbiology and Immunology, Stanford University School of Medicine, Stanford, CA

**The influence of the microscope excitation intensity level on the observed hemi-fusion wait times and extents**

Previous work has demonstrated that single virus fusion measurements can be influenced by the light intensity of the microscope illumination (1). We wished to rule out this artifact for our dengue virus assay.

To test the influence of the excitation intensity, we compared results from our single VLP fusion assay at pH 5 collected using using microscope intensity 10/999 with data collected at intensity 20/999 (**Figure 1**), where the intensity level is as specified by the manufacturer (Lumencor). We observed that the wait time distributions and extents at intensity = 10/999 and 20/999 overlapped within error. However, while the wait time distributions and extents of hemi-fusion were not substantially different between Int = 10/999 and 20/999, we did observe greater evidence of photobleaching at Int 20/999 (sloping downward intensity traces of VLPs over time, not shown). Therefore, we opted to use Int = 10/999 for the fusion data collected herein. We also note that intensities much below 10/999 yielded poor S/N ratios, making it difficult to effectively analyze all particles and fusion events.

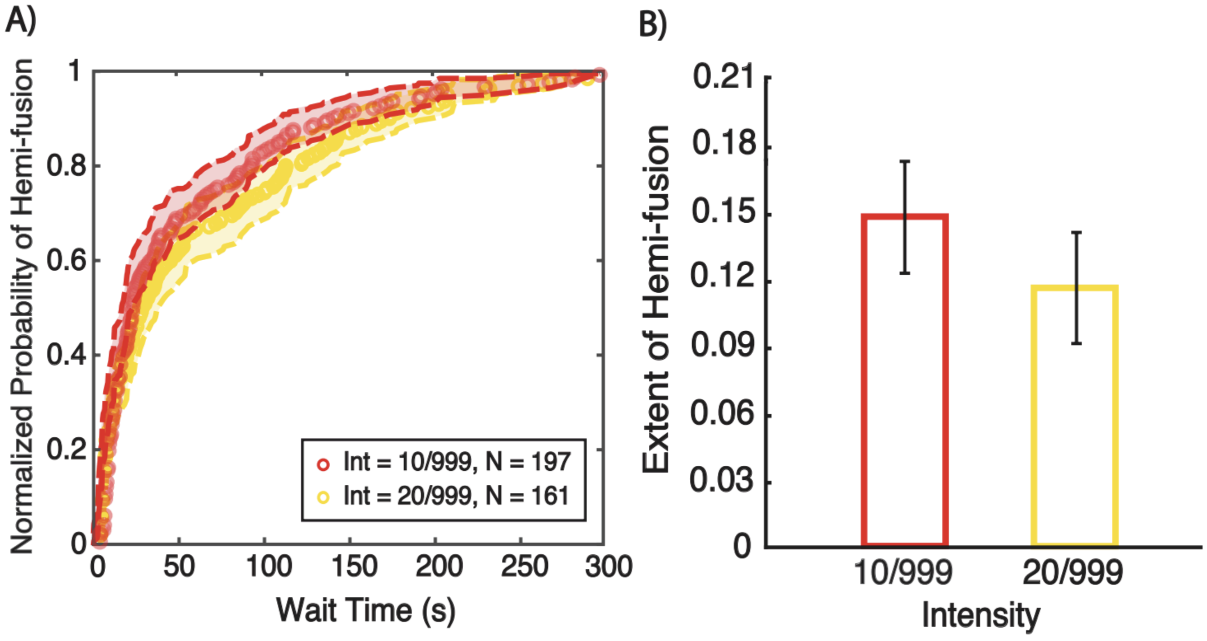


***Figure S1. Influence of microscope excitation intensity level on single VLP hemi-fusion measurements.*** *Single VLP hemi-fusion experiments at pH 5 were monitored using different levels of microscope excitation intensity from the Lumencor Spectra III LED Light Engine (10/999 or 20/999, where the intensity level is as specified by the manufacturer). (A) shows the distributions of observed wait times, displayed as cumulative distribution functions (CDFs). CDFs were normalized to the highest value for each data set. Shaded regions are the 95% confidence intervals determined by bootstrap resampling (NumBootstraps = 10000). (B) shows the extents of hemi-fusion at the different pH values. Extent was measured as the fraction of*

*observed VLPs in the fusion video that underwent hemi-fusion during the 5 min following pH drop. Values shown are the mean ± 95% confidence intervals determined by bootstrap resampling (NumBootstraps = 10000).*

**Determination of the optimal DNA/VLP ratio**

In our single VLP hemi-fusion assay, DNA-lipids were added to VLPs at a set ratio (e.g. 10 DNA/VLP), see Materials and Methods. This DNA/VLP ratio is an important parameter to fine-tune in the assay. If the ratio is too low, very little binding will be observed. If the ratio is too high, the DNA-lipid may "gum up the works", interfering with the hemi-fusion process. To determine the optimal DNA/VLP ratio, we conducted our single VLP hemi-fusion assay at pH 5 using VLPs with ratios of 1 to 200 DNA/VLP, measuring the extent of hemi-fusion at 5 minutes. We observed that the optimal ratio was 10 DNA/VLP, producing the highest extent. Above this ratio, the extent decreased, suggesting that the DNA-lipid interfered with the hemi-fusion process to some extent. At 1 DNA/VLP, substantially less binding was observed (not shown), as expected. The extent was also lower, presumably because a higher fraction of the particles being observed were non-specifically bound. Therefore, we used 10 DNA/VLP in the experiments throughout this report.

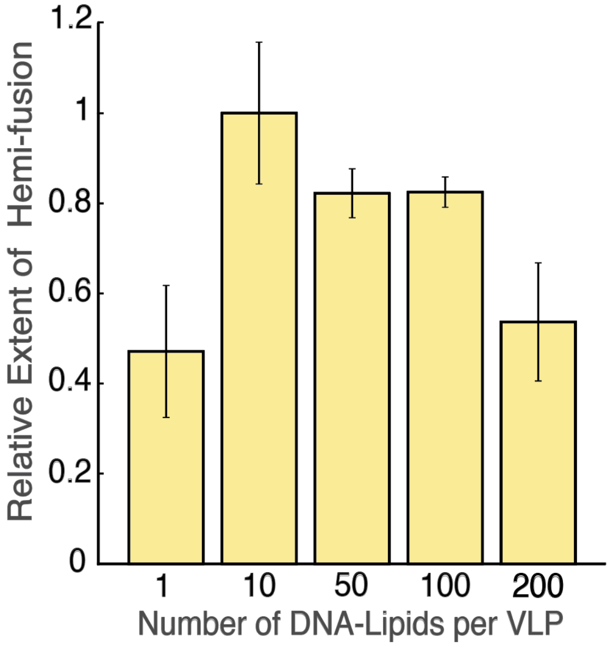


***Figure S2. The dependence of the extent of hemi-fusion on the number of DNA-lipids added per VLP.*** *Single VLP hemi-fusion data was collected at pH 5 using VLPs prepared with different ratios of DNA-lipid per VLP, ranging from 1 to 200 DNA/VLP. Extent was measured as the fraction of observed VLPs that underwent hemi-fusion after 5 min following pH drop.The extents at each ratio are shown, calculated relative to 10 DNA/VLP. Values shown are the mean ± standard deviation of 3 different image regions within each sample.*

**Before/After Fusion Analysis for Measurement of Hemi-fusion Extent**

Extent is defined as the total number of viruses (or VLPs) that underwent hemi-fusion in some time period divided by the total number observed. In previous reports, the extent was measured directly from the fusion video (examples in Refs (2–6)), with viruses that underwent fusion determined by change point analysis of the fluorescence intensity trace (see Materials and Methods and Figure 1b in main text for example trace). Here, we call this the "Traditional" approach. In general, this approach works well, however there are two limitations. First, only one fusion video can be captured per flow cell (because the pH change occurs across the entire flow cell), which limits the total number of viruses that can be observed per experiment. As the extent becomes small (such as at high pH values), this can lead to inaccurate determinations of the true extent as small statistical fluctuations in the number of hemi-fusion events can have an outsized influence. Second, hemi-fusion video micrographs are collected at reasonably low light intensities in order to reduce photobleaching during the measurement (see **Figure S1**). This enables accurate measurement of hemi-fusion wait times, but in our experience can add additional uncertainty to the determination of hemi-fusion extent (number of VLPs hemi-fused/total number of VLPs bound) because dim particles are often not included in the analysis.

Therefore, in this report we developed an alternative method (called the "Before/After" method) to measure the extent of hemi-fusion. In the Before/After method, "before" images are taken in several locations around the flow cell prior to pH drop. These images are taken at a higher light intensity (int = 25/999 on the Spectra III LED Light Engine), and the locations are saved in the microscope software. The pH drop is then carried out by buffer exchange as usual and the flow cell is incubated for the desired time period (usually 5 minutes), after which time the same locations are imaged again in rapid succession using a motorized microscope stage.

To analyze this data, each particle is algorithmically found in the before image and then again in the after image using custom Matlab scripts, and the difference in fluorescence intensity between the two images is calculated. Particles whose fluorescence intensity change is greater than a threshold cutoff are determined to have undergone hemi-fusion during the incubation period. From this, the extent can then be calculated as defined above. The advantage of this approach is that a much larger number of particles can be analyzed in a single experiment, and imaged at higher light intensities, leading to more robust statistics, especially for conditions where the extent is inherently low (such as at high pH values).

For this method to work well, it is essential to determine an appropriate threshold cutoff value. Naïvely, one might assume that an intensity change > 0 would be appropriate, however the process of returning to a previously imaged location and re-focusing will inherently give a distribution of intensity changes, as the re-focusing will not be exact. Therefore, to determine an appropriate threshold cutoff value, we collected before/after data for VLPs that were adhered nonspecifically to a glass coverslip, but otherwise treated the same as a typical hemi-fusion assay run at pH = 5. The intensity change for each particle was collected, and a distribution was constructed (see **Figure S3a**). The threshold was the set as the intensity change that gave a false positive rate of <2%, which in our data was 0.4. This threshold was then applied to all before/after data collected. An example of an intensity change histogram for hemi-fusion data collected at pH 5 is shown for comparison (**Figure S3b**).

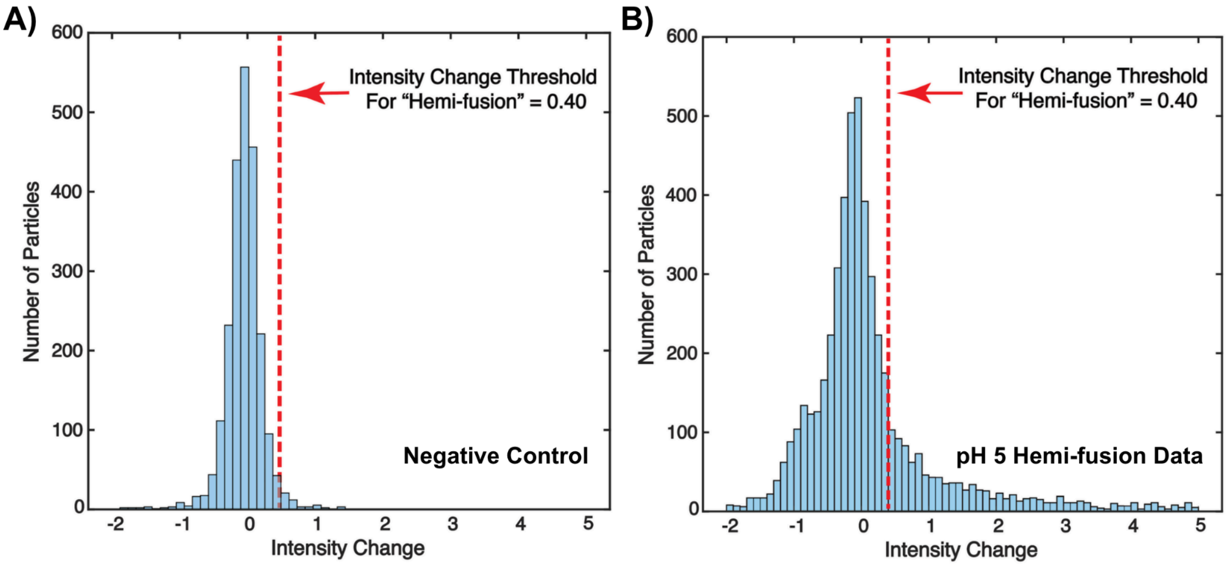


***Figure S3. Histogram of VLP intensity changes in before/after method to measure hemi-fusion extent.*** *(A) "Negative control" shows the histogram of VLP intensity changes in a set of "Before" and "After" images taken of VLPs that were non-specifically adhered to a clean glass coverslip inside a PDMS flow cell and exposed to pH 5 buffer for 5 minutes. This data was used to determine the threshold intensity change of 0.4 (red dashed line), with a false positive rate of <2%. Number of particles analyzed = 2093. (B) "pH 5 Hemi-fusion Data" shows an example histogram of VLP intensity changes in a set of "Before" and "After" images taken in a single VLP hemi-fusion experiment, which was collected at pH 5. Number of particles analyzed = 5517.*

To validate this new approach, we compared our Before/After method of analysis to the Traditional method for a limited data set (a subset of the single VLP hemi-fusion data collected at pH values ranging from 5 to 6.25). We observed good agreement between the 2 approaches, and the trend of decreasing extent with increasing pH was more easily visualized.

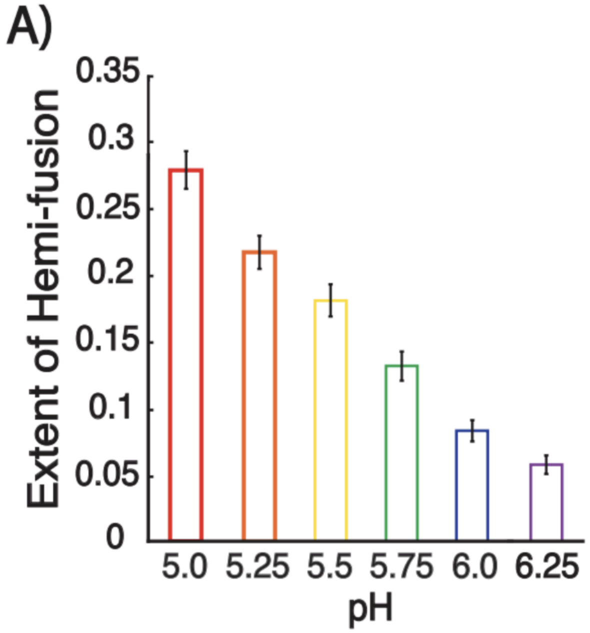


***Figure S4. Comparison of methods to measure hemi-fusion extent.*** *Single VLP hemi-fusion experiments were run at differing pH values, ranging from 5 to 6.25. (A) shows the extents of hemi-fusion at the different pH values as measured using the "Before/After" method – comparing the intensities of the particles in several regions throughout the sample before pH drop with 5 minutes after pH drop. (B) shows the extents of hemi-fusion at the different pH values as measured using the "Traditional" method – analyzing particles in a fusion video collected in a single region of the sample, and identifying hemi-fusion events based on trace analysis. In both cases, extent was defined as the fraction of observed VLPs that underwent hemi-fusion after 5 min following pH drop. Values shown are the mean ± 95% confidence intervals determined by bootstrap resampling (NumBootstraps = 10000).*

**Supporting References (all are cited in the main text as well)**


1. Rawle, R.J., A.M. Villamil Giraldo, S.G. Boxer, and P.M. Kasson. 2019. Detecting and Controlling Dye Effects in Single-Virus Fusion Experiments. *Biophys. J.* 117:445–452.
2. Rawle, R.J., E.R. Webster, M. Jelen, P.M. Kasson, and S.G. Boxer. 2018. pH Dependence of Zika Membrane Fusion Kinetics Reveals an Off-Pathway State. *ACS Cent. Sci.* 4:1503–1510.
3. Cervantes, M., T. Hess, G.G. Morbioli, A. Sengar, and P.M. Kasson. 2023. The ACE2 receptor accelerates but is not biochemically required for SARS-CoV-2 membrane fusion. *Chem. Sci.* 14:6997–7004.
4. Ivanovic, T., J.L. Choi, S.P. Whelan, A.M. van Oijen, and S.C. Harrison. 2013. Influenza-virus membrane fusion by cooperative fold-back of stochastically induced hemagglutinin intermediates. *eLife*. 2:e00333.
5. Chao, L.H., D.E. Klein, A.G. Schmidt, J.M. Peña, and S.C. Harrison. 2014. Sequential conformational rearrangements in flavivirus membrane fusion. *eLife*. 3:e04389.
6. Liu, K.N., and S.G. Boxer. 2020. Target Membrane Cholesterol Modulates Single Influenza Virus Membrane Fusion Efficiency but Not Rate. *Biophys. J.* S000634952030271X.
